# Rapid biomechanical imaging at low irradiation level via dual line-scanning Brillouin microscopy

**DOI:** 10.1101/2022.04.25.489096

**Authors:** Jitao Zhang, Milos Nikolic, Kandice Tanner, Giuliano Scarcelli

## Abstract

Brillouin microscopy is a promising all-optical technique for biomechanics but is limited by slow acquisition speed and/or large irradiation doses. Here, we introduce multiplexed Brillouin microscopy that overcomes both these limits by over one order of magnitude with selective illumination and single-shot analysis of hundreds of points along the incident beam axis. We demonstrate the enabling capabilities of this method probing rapid response to perturbations and long-term mechanical evolution of tumor spheroids.

## Main Text

The biomechanical properties and interactions of cells and tissues are critically involved in many biological functions from cell behavior to tissue-level processes^1, 2^. As a result, many techniques have been developed in the past decades to quantify the mechanical properties of biologically relevant materials^3-6^. Among these, Brillouin optical microscopy has recently emerged as an attractive option due to its ability to quantify the mechanical properties of material with sub-micrometer three-dimensional (3D) resolution and without contact or labels^7-10^. Thus, Brillouin microscopy can provide mechanical measurements when traditional methods cannot be used, for example because no physical access can be gained to the region of interest such as in tissue^11, 12^, 3D microenvironments^13-15^ or microfluidic channels^16^.

Spontaneous Brillouin scattering arises from the interaction of light with thermal acoustic phonons within a material resulting in scattered light^17^. The Brillouin frequency shift, i.e., the difference in frequency between incident and scattered light (∼0.01 nm), is related to the longitudinal elastic modulus and thus gives access to the local mechanical properties of material^18^. Despite tremendous progress in confocal Brillouin microscopy over the past fifteen years, slow acquisition speed (20-200 ms per spectrum) remains the major limiting factor towards the widespread implementation of Brillouin technology in biomedicine^19^. To overcome the speed limitation, stimulated Brillouin scattering has been recently proposed,^20, 21^ where acoustic photons are driven by resonant pump-probe interaction so that much stronger Brillouin signal is generated. However, stimulated Brillouin spectrometers have not reduced acquisition time below 20 ms in biological samples due to sub-optimal continuous wave operation and less efficient light detection. Importantly, both spontaneous and stimulated Brillouin microscopy are based on point-scanning to cover the 3D volume of a sample leading to slow acquisition times and high irradiation doses given the redundant illumination of out-of-focus voxels. This challenge has made Brillouin mapping often impractical for applications in multicellular organisms or tissue engineering where biomechanics is of crucial relevance^22, 23^.

An efficient solution to the speed/photodamage challenge lies in multiplexing. In other fields for example, light-sheet fluorescence microscopy has improved acquisition speed and reduced photodamage compared with confocal fluorescence microscopy by selective illumination and multiplexed detection^24^. Here, for the first time, we developed a viable multiplexing solution for Brillouin spectroscopy/microscopy via selective line-scanning Brillouin microscopy (LSBM) with simultaneous imaging and single-shot spectral analysis of hundreds of points. The setup features dual-line illumination with two counterpropagating beams (**Figure 1a & 1b, Supplementary Fig. 1**), which enables light-efficient multiplexing along the illumination beam axis and orthogonal collection free of artifacts due to heterogeneous refractive index distribution (**Supplementary note 1, Supplementary Fig. 2**). To enable single-shot Brillouin spectral analysis of the multiple illuminated points, we used the perpendicular direction to the dispersion axis of an etalon interferometer^25^ (based on a virtually-imaged-phased-array (VIPA) etalon) for spatial multiplexing. To further minimize absorption-induced phototoxicity, we used a narrowband infrared laser and cleaned its spectrum to suppress side modes and amplified spontaneous emission (**Supplementary Fig.1, Methods**). Using such laser also enabled additional 40 dB rejection of non-Brillouin light with a heated Rb gas cell by locking the laser frequency to a Rb absorption line. Overall, the multiplexed Brillouin spectrometer reaches all the required specifications for biomechanical analysis of biological samples (**Supplementary Fig. 3**): spectral extinction >70 dB for shot-noise operation; spectral precision of <10 MHz, leading to 0.28% relative precision needed to detect subtle changes; and equivalent acquisition of 1 ms per pixel in complex biological samples, i.e. over one order of magnitude faster than state-of-the-art spontaneous confocal or stimulated setups.

**Figure 1.**
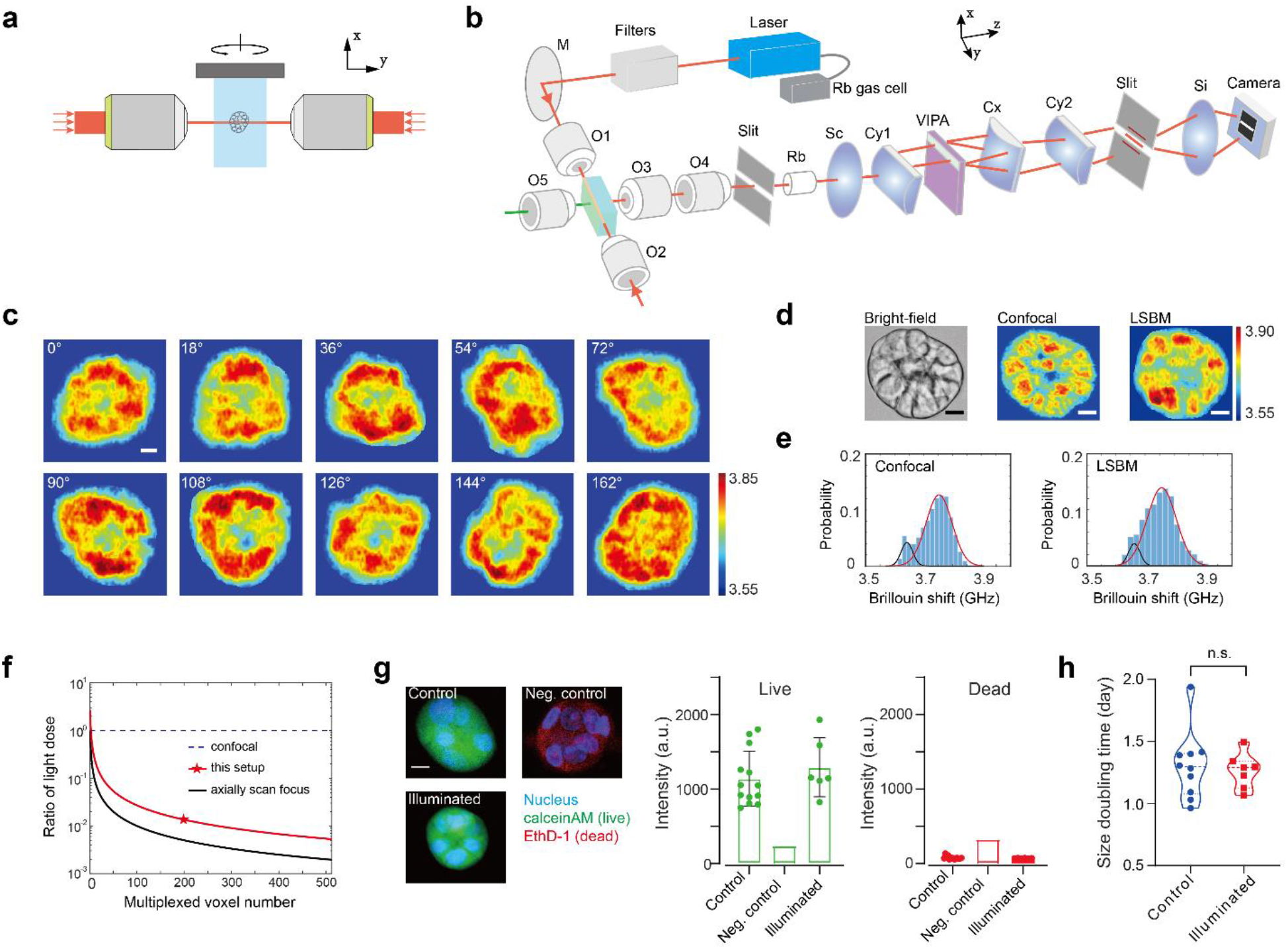
Design and validation of the LSBM. **a**, Schematic of dual-line illumination configuration. **b**, Optical setup of the instrument. The laser beam, whose frequency is locked to the absorption line of Rubidium gas, is spectrally cleaned by filters and focused into the sample from two sides. A home-built bright-field microscope is used for image guidance, and the Brillouin signals are collected by the multiplexed spectrometer. M, mirror; O1-O5, objective lenses; Rb, Rubidium gas chamber; Sc, Si, spherical lenses; Cy1, Cy2, Cx: cylindrical lenses. **c**, 3D Brillouin mapping of a spheroid by rotating it along x axis. Labels indicate the relative azimuth angle in y-z plane. **d**, Brillouin images of a spheroid measured by confocal Brillouin modality and the LSBM. **e**, Histogram distributions of the Brillouin shifts of the scanned sections by confocal Brillouin and LSBM. Solid lines are fitting results by Gaussian profile. The peak locations are 3.64 GHz and 3.75 GHz for confocal image and 3.66 GHz and 3.75 GHz for line-scanning image, respectively. **f**, Induced light dose of the LSBM setup compared to confocal configuration for 3D Brillouin imaging. **g**, Live-dead assay indicates the light dose induced by the LSBM does not affect the viability of the sample. Control group, n=13; Illuminated group, n=6. **h**, The proliferation rate of the illuminated group (n=7) does not show significant difference compared with the control group (n=10). n.s., not statistically significant. Statistical significance is determined by performing unpaired t-test. Scale bar, 10 µm. Color bar represents Brillouin shift with a unit of GHz.

To enable imaging simultaneous to spectral analysis, the multiplexed etalon interferometer was aligned in the infinity space of a 2.5x imaging system, and the dual illumination lines were conjugated with a slit (which thus provided additional confocal sectioning) and the entrance window of the etalon. Moreover, the image of the dual illumination beam and the Brillouin spectra were formed at the same plane using orthogonal cylindrical lenses (**Figure 1b**). To acquire a 2D Brillouin image, the sample was scanned along the *x*-axis. To reconstruct a 3D image, multiple 2D Brillouin images were obtained by rotating the spheroid in the *y*-*z* plane (**Figure 1c**). Overall, using illumination NA of 0.1 and collection NA of 0.3, the LSBM has a spatial resolution of 1.5 µm×1.5 µm×4.8 µm. For validation, we compared our LSBM setup with a standard confocal Brillouin microscope on the same sample (**Figure 1d**). Although the co-registration of images cannot be exact due to fundamental (voxel size of LSBM is ∼2x smaller) and practical (the sample is manually transferred between instruments) reasons, the histogram distributions of the images indicate LSBM successfully captured the same mechanical features of a spheroid as the confocal Brillouin microscope (**Figure 1e**).

Next, we demonstrated that the speed improvement of our LSBM configuration is accompanied by a marked decrease in irradiation dose. Since spontaneous Brillouin scattering operates in non-depleted-pump regime, and thanks to the dual-line illumination (**Supplementary note 1**), we could implement “on-axis” multiplexing, i.e. we used the illumination beam propagation axis (here noted as *y*-axis) as the multiplexing direction. This effectively eliminates the out-of-focus redundant illumination of point-scanning or off-axis multiplexing configurations thus resulting in the total light dose being reduced to 7.4% of an equivalent confocal setup (**Figure 1f**, red star) for 3D imaging (**Supplementary note 2**). The light dose could be further reduced by 5 times (**Figure 1f**, black curve) using a tunable lens rather than a fixed objective for illumination. We next checked the viability and the proliferation rate of the spheroids after illumination: we verified that LSBM’s light dose not only does not introduce any photodamage (**Figure 1g**), but, more importantly, it does not affect spheroids’ growth (**Figure 1h, Supplementary Fig. 4**), a clear demonstration of the non-perturbative nature of our configuration. In summary, we have demonstrated that LSBM provides more than one order of magnitude improvement in both acquisition speed and light dose, thus enabling Brillouin imaging for both rapid response and long-term biomechanical studies.

Tumor organoids are a popular and effective *in vitro* model to study tumor etiology, progression, and drug sensitivity^26, 27^ in which mechanical properties are recognized to be important but have remained elusive^28^; thus, they offer a valuable system to field-test the features of multiplexed Brillouin technology. The improved acquisition speed of the LSBM allowed us to monitor the quick mechanical response of tumor spheroids to external stimuli such as osmotic stress. Osmotic stress is an important regulator of cell function in mechanobiology and affects gene expression and metabolic activity^29, 30^. We subjected tumor spheroids to osmotic shocks and tracked their Brillouin shift every minute (**Figure 2a-2c**). We observed that the Brillouin shift of the multicellular spheroids increased under hyperosmotic shock and decreased under hypoosmotic shock (**Figure 2d, Supplementary Fig. 5**) as expected^31^. Beyond this, Brillouin imaging ability revealed that the internal mechanical heterogeneity had dramatically different response to hypoosmotic shocks, where spheroids became spatially uniform, compared to hyperosmotic shocks, where spheroids maintained similar spatial heterogeneity as control samples (**Figure 2e**).

**Figure 2.**
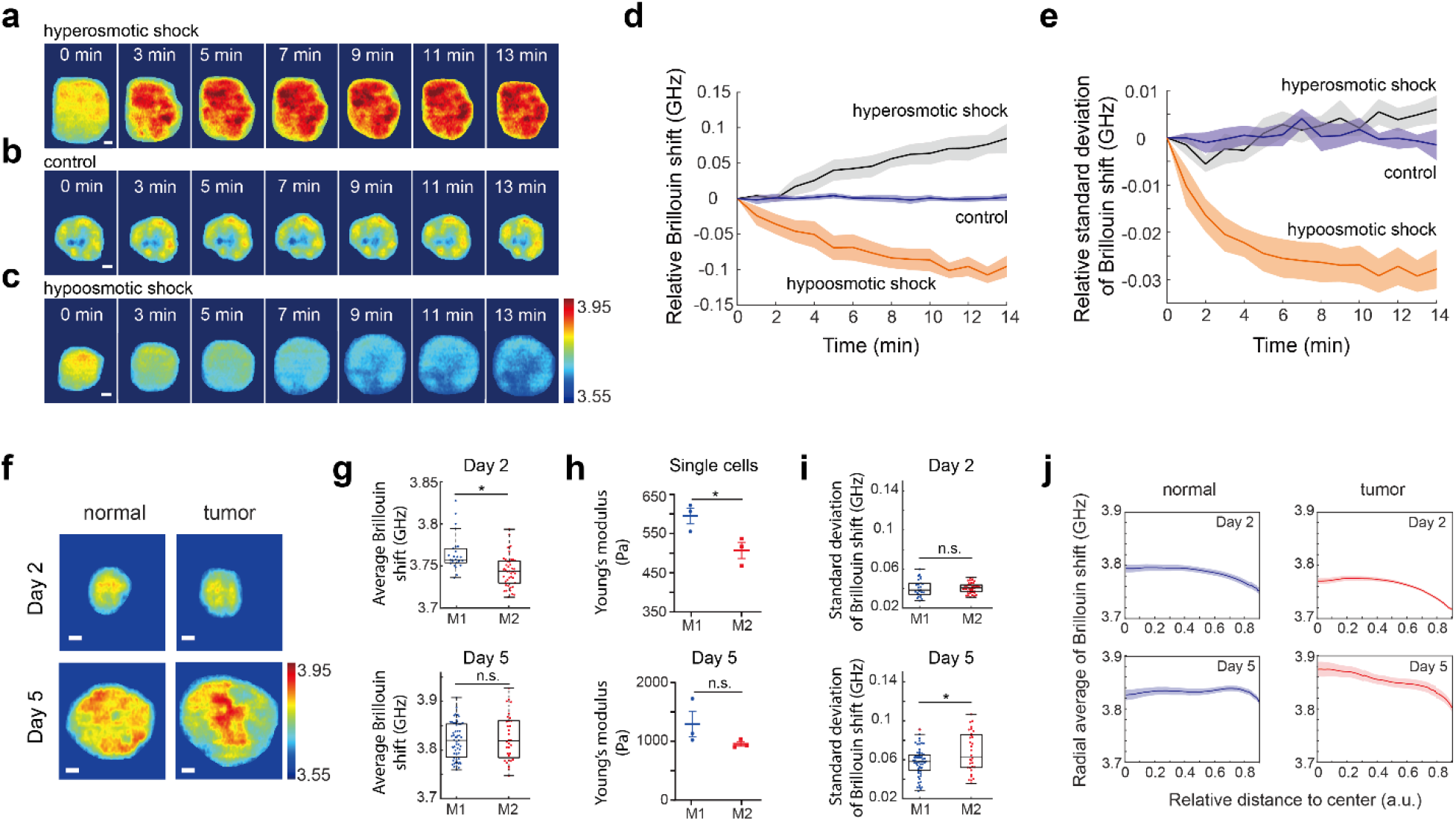
Mechanical response of the spheroids to external perturbations and long-term mechanical evolution. **a**-**c**, Representative time-lapse Brillouin images of the spheroid under conditions of hyperosmotic shock (**a**), no shock (**b**), and hypoosmotic shock (**c**). Scale bar, 10 µm. **d, e**, Temporal change of the averaged Brillouin shift (**d**) and the standard deviation (**e**) of spheroids in hyperosmotic shock group (n=5), control group (n=6), and hypoosmotic shock group (n=5). **f**, Representative Brillouin images of normal (M1) and tumor spheroid (M2) on Day 2 and Day 5. **g**, On Day 2, M1 (n=26) has higher Brillouin shift than M2 (n=42). \**p*=0.0023. While on Day 5, there is no significant difference between M1 (n=59) and M2 (n=32). Statistical significance is determined by performing unpaired t-test. **h**, The results of AFM experiment show similar trend of the mechanical evolution as Brillouin experiments. Data points show the mean of all independent repeats. \**p*=0.0379. Statistical significance is determined by performing unpaired t-test with all repeats. **i**, M1 and M2 show no difference on standard deviation of the Brillouin shift on Day 2 but are significantly different on Day 5. \**p*=0.0074. Statistical significance is determined by performing unpaired t-test. **j**, Radial averages of the Brillouin shifts of normal and tumor spheroids on Day 2 and Day 5. Error bars represent ±s.e.m. Color bar represents Brillouin shift with a unit of GHz.

Within tumor organoids, tumor cells interact via mechanical coupling with nearest cellular neighbors and the extracellular matrix (ECM). Recent work has revealed that tumor cells show different mechanical properties as a function of the ECM milieu compared to that of their normal counterparts^32-34^. However, how and what drives this mechanical coupling across multicellular structures is less understood. This is an important open question on how tumor cells adapt to *de novo* mechanical environments along the metastatic cascade. Here, using spheroids derived from breast cancer isogenic cancer progression lines that mimic different stages of malignancy, we demonstrated that LBSM enables observing unique mechanical features during growth. We compared the mechanical evolution of healthy tissue spheroids (from healthy epithelial cell line (M1: MCF10A)) with the mechanical properties of tumor-like tissue spheroids (from tumor cell line (M2: MCF10AT1k.cl2)). At single cell level, healthy mammary cells have higher modulus than the corresponding precancerous or cancerous ones^35^; here we asked if such relation was conserved upon spheroid growth. We measured spheroids at early (Day 2) and later (Day 5) culturing stage (**Figure 2f, Supplementary Fig. 6**). At day 2, tumor spheroids showed lower Brillouin shift than healthy spheroids, mirroring the single cell behavior. However, as spheroids grew, this mechanical difference vanished (**Figure 2g**), a new observation which we validated with atomic force microscopy (AFM) (**Figure 2h, Supplementary Fig. 7**): single healthy cells are significantly stiffer than cancerous ones, but as spheroids grow, the stiffness of tumor spheroids increases more rapidly so that by day 5 healthy and tumor spheroids have similar stiffness. Beyond average values, LSBM enables to investigate the spatial behavior of the mechanical evolution of multicellular spheroids. We observed that tumor spheroids showed higher mechanical heterogeneity than normal ones while growing (**Figure 2i**). To further understand this feature, we extracted the radial average of the Brillouin shift (**Figure 2j**). We found both normal and tumor spheroids had similar radial mechanical change at early stages. Instead, at later stages, the tumor population showed much steeper radial gradient than the normal spheroids, suggesting that the rapid change in mechanical properties is driven by the core of the spheroids. Since tumor progression is often accompanied by altered tissue biomechanics, the LSBM can serve as a unique platform to dissect the role of mechanical regulation in tumorigenesis.

In summary, we have demonstrated rapid 3D mechanical mapping of biological samples at micron-scale resolution and low non-perturbative light dose thanks to dual line-scanning Brillouin microscopy featuring simultaneous imaging and single-shot spectral analysis of hundreds of points along the optical axis. Several improvements of the technique could be implemented from advances known in light-sheet fluorescence microscopy: e.g. the efficiency of the technique could be improved by axially scanning the focus of the illumination beam; analysis of different sample types as well as easier sample preparation could be obtained with different mounting configurations. We validated the technique against confocal Brillouin microscopy verifying the equivalence of extracted information as well as the advantageous instrument performances in terms of speed, precision and light dose. We demonstrated LSBM enables to detect biologically relevant mechanical changes in multicellular organisms such as tumor spheroids for both short time studies and long-term culture, limited only by the penetration depth of the illumination beam due to spheroid turbidity. Importantly, in this demonstration, we discovered a previously unreported robust stiffening behavior in the evolution of growing tumor spheroids which we validated against gold standard AFM. These capabilities show LSBM is a promising method to study biomechanical processes in developmental biology, tissue engineering and organ-on-a-chip applications.

## Acknowledgments

We thank Dr. Giulia Zanini for help with photodamage experiment. This work was supported by National Science Foundation (CMMI-1929412), the National Institutes of Health (R33CA204582, K25HD097288), the Intramural Research Program of the National Cancer Institute, and American Cancer Society Institutional Research Grant (1816016).

## Author Contributions

J.Z. and G.S. conceived the project. J.Z., M.N., K.T., and G.S. devised the research plan. J.Z. developed the instrument and performed the experiments. M.N. developed the spheroid protocols, performed photodamage experiment and AFM measurement. J.Z. and G.S. wrote the manuscript with input from all other authors.

## Online methods

### Dual line-scanning Brillouin microscope

The light source is a continuous wave tunable diode laser (DL pro, Toptica) with a central wavelength of 780.24 nm and a linewidth of less than 0.3 MHz. During operation, the laser frequency was locked to the absorption line of the Rubidium gas (Rb-85). The light source was coupled into an optical amplifier (BoosTA, Toptica) to acquire an output power as high as 3 watts. The output beam was delivered to the optical setup via an optical fiber and a collimator (PAF2A-11B, Thorlabs). The laser spectrum was cleaned by using two ultra-narrowband Bragg filters (BP-780, OptiGrate) and a Fabry-Perot etalon (Light Machinery) with a free spectrum range (FSR) of 15 GHz. The laser beam was then expanded to about 9.5 mm and sent into an objective lens (4x/0.1NA, Olympus) to create a loose beam line for illumination. For dual-line illumination, a flip mirror was used to guide the beam to create a second identical beam line that is propagating against the first one (**Supplementary Fig.1**). The alignment of the two beams was achieved with the help of the live side image taken by a home-built imaging system, which includes an objective lens (10x/0.25, Olympus), a tube lens (Thorlabs), and a sCMOS camera (Neo, Andor). The beam lines were superposed within a 1-cm sized cuvette (21-200-255, Fisher Scientific), which was filled with medium and used as the sample holder.

The Brillouin signals generated on the illumination beam line were collected with 90-degree geometry by a multiplexed Brillouin spectrometer (**Fig.1b**). The beam line was first imaged onto the first slit (VA100, Thorlabs) by a pair of objective lenses (20x/0.4NA, 4x/0.1NA, Olympus), which yielded the effective NA of ∼0.3 for the collection path. The image of the line was then collimated by a spherical lens (f = 400 mm, Thorlabs) and coupled into the entrance window of the VIPA etalon (FSR=10 GHz, Light Machinery) by a cylindrical lens (f = 200 mm, Thorlabs). After the VIPA etalon, the spectrum of the beam line was projected onto the second slit (VA100, Thorlabs) by a combination of two cylindrical lenses (f_1_ = 1000 mm, f_2_ = 400 mm, Thorlabs). The Brillouin spectral pattern was then reimaged by a doublets pair (MAP1040100-B, Thorlabs) and recorded by a EMCCD (iXon Ultra 897, Andor). To improve the spectral extinction, a 150-mm Rb gas cell (TG-ABRB-I85-Q, Precision Glassblowing) was placed between the first slit and the collimation lens. When heated up to around 65 °C, the gas cell can provide spectral extinction of ∼ 40 dB.

A customized four-dimensional stage (three translational movements and one rotational movement) was integrated with the LSBM setup for scanning the sample. We developed a LabVIEW-based program for setup operation and data acquisition. For spheroid imaging, we used the input laser power of ∼370 mW and set the exposure time of the EMCCD as 200 ms. Since 200 pixels were acquired simultaneously, the equivalent acquisition time for each pixel is 1 ms. The spectral analysis was conducted in MATLAB (R2021b). The Brillouin shift was retrieved by fitting the spectrum with Lorentzian profiles. We used standard materials including water and methanol for the calibration of the spectrometer.

For dual-line illumination imaging, the ultimate image was obtained by image fusion process using MATLAB. To do this, we adapted a procedure established in light-sheet microscopy^36^. For each sample, we first took two Brillouin images under single-line illumination condition. We then aligned the single-illumination images based on the profile features and the bright-field image of the sample. Next, we spatially overlapped two images. For each pixel, we selected the value with larger Brillouin shift, and inserted it into a new fused data set. The final image was obtained by cropping the region of interest of the fused data set.

### Confocal Brillouin microscope

A standard confocal Brillouin microscope was used for validation. The details of the instrument can be found in previous report^10^. Briefly, a 660-nm continuous wave laser (Torus, Laser Quantum) with ∼10 mW was used as the light source. The add-on Brillouin module was integrated with a commercial inverted confocal microscope (IX81, Olympus) for conducting 2D/3D mapping. A two-stage VIPA based Brillouin spectrometer was used for acquiring Brillouin signal. An objective lens with NA=0.4 (LMPLFLN20x, Olympus) was used for spheroid imaging.

### Cell culture

MCF10A (M1) and MCF10AT1k.cl2 (M2) cells were received from the Barbara Ann Karmanos Cancer Institute (Detroit, MI, USA). They were cultured according to the supplier protocol, and early passages were cryopreserved and thawed as needed. All cells used were passage number <15. Cells were cultured in T25 flasks at 37°C, 5% CO_2_, in complete medium consisting of DMEM/F12 (11330-032, Thermo Fisher Scientific), 5% horse serum (16050-122, Thermo Fisher Scientific), 5 ng/ml EGF (AF-100-15-1MG, Peprotech), 0.5 mg/ml Hydrocortisone (H0888-1G, Sigma-Aldrich), 100 ng/ml Cholera toxin (C8052-2mg, Sigma-Aldrich), 10 µg/ml insulin (I1882-100MG, Sigma-Aldrich) and 1x penicillin/streptomycin solution (15070-063, Thermo Fisher Scientific). Cells were passaged at around 80% confluency using 0.05% trypsin (25-052-Cl, Corning) for 5 min. After treatment with trypsin cells were centrifuged at 150g for 5 min, resuspended in fresh medium, and seeded in a new T25 flask. Cell medium was changed every 2-3 days.

### Spheroid morphogenesis assay in 3D on-top Matrigel culture

Previously established protocols to form acinus-like spheroids of M1 and M2 cells in 3D on-top culture were followed^37,38^. Spheroids were cultured in 2-well glass bottom imaging slides (Ibidi, 80287). 200 µl of ice-cold Matrigel (Corning, 356231) was added to a well of an imaging slide that was previously chilled on ice. After spreading the Matrigel evenly on the bottom of the well with a pipette tip, the slide was placed in the incubator for 30 min for the layer of Matrigel to solidify. Cells were harvested with trypsin from the flask as described above. After centrifugation, the supernatant was aspirated, fresh medium was added, and cells were mixed thoroughly with a pipette to ensure single cell suspension. Next, cells were counted using a hemocytometer. 20,000 cells were carefully mixed in 500 µl of the assay medium. Assay medium is identical to the complete medium with the exception that it contains only 2% horse serum. After that, 500 µl of the cell solution was added on top of the Matrigel layer in a well of a 2-well slide. The slide was placed in the incubator for 30 min to allow the cells to settle on top of the Matrigel layer. After that, 500 µl of assay medium containing 10% Matrigel was carefully pipetted on top of the cells in the well. This yields a final concentration of 5% Matrigel in assay medium. Acinus-like spheroids will form by day 5 of growth. The medium in the well was changed every 2 days by carefully aspirating the medium and replacing it with 1 ml solution of 5% Matrigel in assay medium.

### Sample preparation for Brillouin imaging

Spheroid sample was collected from the culture dish by gently dislodging the spheroids from the Matrigel in the well with a pipette tip and transferring them to an Eppendorf tube with a pipette. After 5-10 seconds centrifuging with tabletop centrifuge (VWR, Mini Centrifuge), most of the supernatant was removed, and the remaining spheroids at the bottom of the Eppendorf tube were resuspended into 100 µL 1% agarose phosphate buffer saline (PBS) solution which was previously warmed to ∼37°C. The solution was carefully pipetted a few times to allow an even distribution of the spheroids. The solution was then injected into a FEP tube with an internal diameter of 1/16 inches (Cole-Parmer). After allowing 5 min for agarose gelation at room temperature, the agarose gel cylinder which contains embedded spheroids was partially pushed out from the tubing using a pipette tip. For LSBM experiment, the tubing was mounted onto the 4D stage, and the gel cylinder was immersed into the PBS solution of the cuvette. For confocal Brillouin experiment, the tubing along with the gel cylinder was placed into a glass bottom petri dish containing PBS solution.

To perform the osmotic shock experiments we replaced the medium in the cuvette with a hyperosmotic solution (500 mM sucrose), or hypoosmotic solution (25% PBS, 75% dH_2_O). We immersed the spheroids embedded in the agarose gel directly into the solution and proceeded to acquire a time series of Brillouin shift maps.

### Photodamage experiment

On day 0, cells were seeded in 2-well slides with 500 µm grid on the bottom (Ibidi) as described above. On day 2, the spheroids in the well (without dislodging them) were placed on the microscope stage, and individual spheroids and their locations on the annotated 500 µm grid were recorded. We illuminated the spheroids with 780 nm light, 370 mW, for 5 minutes, using the same objective which was used to create the illumination beam in the Brillouin line scan experiments (4x, 0.1 NA, Olympus).

To estimate cell viability at the end of the experiment, all spheroids in the well were stained with 2 µM calcein AM, and 5 µM Ethidium Homodimer-1 (Live/Dead Viability/Cytotoxicity kit, ThermoFisher Scientific) in PBS for 45 min. Sample was then washed with PBS and imaged using the confocal microscope (Olympus FV3000) with a 10x, 0.4 NA objective (Olympus, UPLSAPO10X2). Calcein AM fluorescence was observed in the green channel (488 nm excitation, 500-540 nm emission), and Ethidium Homodimer was observed in the red channel (514 nm excitation, 590-690 nm emission). We collected z-stacks (1.4 µm pixel size in x-y plane, 3.9 µm step in z direction). Maximum intensity projection image of each spheroid was thresholded in the green channel and average fluorescence intensity of the thresholded region was calculated for both green and red channels with image processing software Fiji.

To estimate the rate of spheroid growth we tracked spheroids based on their location on the 500 µm annotated grid. We recorded 10x, 0.1 NA brightfield images of the illuminated and control spheroids on each day until day 5. Projected area of the spheroid was manually selected in Fiji.

The area versus day data was fitted using an exponential growth model *A* = *A*_0_*e*^*kt*^ (Prism 8, GraphPad) where A is the spheroid projected area, A_0_ is the initial area, k is the growth rate and t is time in days. The doubling time was calculated as ln(2)/*k*. Difference between doubling times of illuminated and control spheroids was tested using unpaired, two-tailed, t-test with Welch’s correction. (Prism 8, GraphPad).

### Cell and spheroid stiffness measurement using AFM

Cell and spheroid stiffness was measured using the NanoWizard 4a AFM (JPK Instruments). AFM cantilever with 5 µm diameter round tip was used for indentation (Nano and More, CP-qp-CONT-Au-B). The spring constant of the cantilevers was calibrated using the thermal noise method while they were immersed in the cell medium in the dish. For single cell measurement, cells suspended in 2 ml of assay medium were added to the glass bottom dish (World Precision Instruments, FluoroDish) and allowed 2 minutes to settle on the glass bottom. Tip was centered on the cell body and each cell was indented with the maximum force setpoint of 5 nN and with approaching speed of 2µm/s across the height range of 5 µm. The measurement was repeated 9 times in the same location and each indentation curve was recorded. For spheroid measurements on day 5 of growth, the spheroids were physically detached from the Matrigel bottom layer in the well with a pipette tip and transferred to an Eppendorf tube. They were centrifuged for 5-10 seconds using the tabletop centrifuge, and supernatant which may contain chunks of Matrigel was removed. Spheroids were then carefully resuspended in 2 ml of assay medium and added to a glass bottom dish. After they were allowed 2 minutes to settle, AFM tip was centered on each spheroid and indented 9 times in the same location with the maximum force setpoint of 25 nN, with approaching speed of 2 µm/s across the height range of 15 µm. The height of spheroids was measured by approaching the AFM tip to the top of the spheroid using a 1 nN setpoint and recording the tip height. For each spheroid, we also recorded the tip height after approaching the glass bottom near the same spheroid. The AFM indentation curves were analyzed using the JPK data processing software. Indentation curves were fitted using the Hertz/Sneddon model on the 75% of the curve, assuming the Poisson ratio of 0.5 and a spherical indenter with a diameter of 5 µm. For each cell or spheroid, Young’s moduli from all 9 indentation measurements were averaged. At least 5 cells or spheroids were measured in each condition, and each experiment was repeated 3 times. Statistical differences were tested using unpaired, two-tailed, t-test with Welch’s correction. (Prism 8, GraphPad).

## Data availability

The authors declare that all data supporting the findings of this study are available within the paper and its Supplementary note files.

## Code availability

The authors declare that all custom codes of this study are available upon request.

