## Supplementary Information for "Rapid biomechanical imaging at low irradiation level via dual line-scanning Brillouin microscopy"

**Supplementary note 1: Dual-line illumination configuration**

A line-scan configuration in orthogonal geometry, i.e. multiplexed "on-axis" along the illumination beam is highly desirable for rapid detection and low illumination dose (see **Supplementary Note 2**). However, orthogonal microscopy configurations, such as light sheet fluorescence microscopy or on-axis LSBM, carry inherent issues because the illumination and detection paths are affected by the optical property of the sample. Specifically, the index mismatch between sample and medium, the inhomogeneous distribution of the refractive index of the sample and its turbidity will deviate the propagation trajectory of the illumination/detection beams, which causes a decrease of signal intensity and potential distortion of the image. In Brillouin microscopy, this issue also leads to an additional artifact because the value of the Brillouin shift depends on the angle between illumination and collection path, thus a deviation from the orthogonal geometry needs to be carefully considered. To minimize these issues and enable on-axis LSBM, we developed a dual-line illumination configuration, where two beams are counter-propagated and sequentially serve as illumination beams for Brillouin acquisition (**Fig. 1a & 1b, Supplementary Fig. 1**).

The dual line illumination solution comes from both experimental and simulation studies of Brillouin microscopy in orthogonal geometry: we observed that with single-line illumination, there is a noticeable gradient of Brillouin shift along the illumination path due to a deviation from ideal geometry (e.g. **Supplementary Fig. 2a & 2b)**. The deviation from orthogonal geometry is mostly caused by two distinct effects: (A) redirection of the illumination beam path; and (B) redirection of the scattered light path. In case A, given the cylindrical shape of the sample holder and the nearly spherical shape of the spheroids, we avoided artifacts in straightforward manner by aligning the illumination beam with the center of the spheroid. As a result, any beam redirection will be restricted to the illumination plane and the orthogonal scattering geometry is maintained. In case B, artifacts are difficult to avoid with single line illumination. The index mismatch between sample and medium as well as the curved sample geometry create a lensing effect that reduces the angle of the scattered light collected by the spectrometer thus introducing a decreasing gradient in Brillouin shift. This effect could be corrected in post-processing if one collected a 3D map of the sample's refractive index. We instead effectively removed this artifact with two illumination beams counter-propagated and overlapped within the field of view. **Supplementary Fig. 2** exemplifies our procedure. Since the artifact always results in a decreased Brillouin shift along propagation direction, we adapted a procedure developed in light-sheet microscopy[1]: for each sample, we first took Brillouin images under single-line illumination condition (**Supplementary Fig. 2a & 2b**); then, using MATLAB, we optimized co-registration of the single-line illumination images based on the profile features and the bright-field image; next, in the overlapped images, for each pixel we selected the value with larger Brillouin shift to obtain the artifact-free 2D image (**Supplementary Fig. 2c)**.

Interestingly, the dual-line illumination is equipped with a translational degree of freedom along the propagation axis so that with illumination NA of 0.1, the field of view of Brillouin imaging along the multiplexed axis can be varied from ~150 µm to ~300 µm. In experiment, we have measured spheroids with diameter of up to ~100 µm; this limit was set by the penetration depth of the illumination beam due to the turbidity of the spheroids[2].

**Supplementary note 2: Total light dose introduced by confocal vs line-scanning configurations for 3D Brillouin imaging**

The total energy delivered to a sample depends on the illumination intensity $I$, the area of the illumination $A$, and the total acquisition time, i.e. the dwell time $t$ for acquiring one spectrum/voxel times the number of imaged voxels $N_i$ ($i = x, y, z$ represents the pixel number in each dimension).

In confocal Brillouin microscopy, to acquire a 3D image, one usually first collects a 2D section ($N_x \times N_y$ voxels) within the focal plane by point scanning; then moves the focal plane to a different depth in z-direction and collects another 2D section. This strategy is repeated until all the sections ($N_z$) are collected. As a result, the total energy delivered to the sample is:

$$E_{con} = I_{con} \cdot A_{con} \cdot t_{con} \cdot N_x \cdot N_y \cdot N_z \quad (1)$$

The inherent inefficiency of this scanning mechanism is well known: when collecting a voxel in a given plane, the light beam illuminates a large cone of voxels above and below the focal plane.

The line-scanning configuration multiplexes the process of illumination-detection to a line of voxels, thus leading to much quicker image acquisition time. In terms of energy dose, the multiplexing does not necessarily guarantee lower levels because in a linear process, like spontaneous Brillouin scattering, simultaneous acquisition of $K$ voxels generally requires increasing the illumination intensity by a factor $K$ to achieve the same SNR at each voxel. However, since spontaneous Brillouin is a weak scattering process (scattering efficiency is about $10^{-9}$), we can effectively operate in non-depletion pump condition, which enables to multiplex many voxels along the incident optical axis without increasing illumination intensity.

To take full advantage of this effect, we aligned the incident beam line along the y direction and thus multiplexed the scanning by 200-fold (**Figure 1a & 1b**). In this scenario, to obtain a 3D image, the beam line needs to be scanned only once in the x direction, and once in z direction (or equivalently at different angles to obtain the 3D volume in spherical symmetry). As a result, the total light dose delivered to the sample is:

$$E_{ls} = I_{ls} \cdot A_{ls} \cdot t_{ls} \cdot N_x \cdot N_z. \quad (2)$$

The ratio of the light dose introduced by two configurations is

$$R = E_{ls}/E_{con} = \alpha/N_y\ , \quad (3)$$

where $\alpha = I_{ls}/I_{con} \cdot A_{ls}/A_{con} \cdot t_{ls}/t_{con}$. In the ideal scenario, both configurations have similar experimental parameters, i.e. $\alpha = 1$, and the ratio of the light dose will be simply improved by multiplexed number of voxels along the optical axis $N_y$. However, both the illumination areas and the acquisition times per voxel introduce design tradeoffs that need to be evaluated in more depth.

In terms of illumination areas, the two configurations differ because in the confocal epi-detection arrangement the illumination and detection volumes are perfectly overlapped while the 90-degree geometry of line-scanning configuration introduces a mismatch between the size of the incident beam line and the size of the sampled volume. The illumination areas are determined by the numerical apertures (NAs) of the incident beams: $A_{ls}/A_{con} = NA_{con}^2/NA_{ls-ill}^2$, where $NA_{con}$ and

$NA_{ls-ill}$ are the NA of the confocal configuration and the illumination beam of the line-scanning configuration. In our experiment (red line in **Figure 1f**), we created the line scan by focusing a laser beam with a single lens: thus, we used a low illumination NA=0.1 but matched the detection NA (0.3) to the equivalent confocal NA. This lowered the efficiency term of illumination areas, but led to an improved, nearly isotropic, resolution in the line-scan configuration

In terms of acquisition times per voxel, the 90-degree geometry of the line-scanning configuration requires slightly longer collection time because of lower Brillouin geometrical efficiency. The collected Brillouin scattering signal can be written as $\epsilon = I_{ill} \cdot V \cdot \Omega \cdot S$, where $I_{ill}$ is the intensity of the illumination light, $V$ is the interaction volume of the scattering, $\Omega$ is the collected solid angle, and $S$ is the scattering coefficient, which depends on the inherent property of the material and can be considered as a constant here. Thus, the collected scattering signals for confocal and line-scanning configurations are respectively:

$$\epsilon_{con} = I_{con-ill} \cdot S \cdot \pi^2 \cdot 0.61^2 \cdot \lambda^3 / NA_{con}^2 \, , \tag{4}$$

$$\epsilon_{ls} = I_{ls-ill} \cdot S \cdot \pi^2 \cdot 0.61^3 \cdot \lambda^3 / NA_{ls-ill} \, . \tag{5}$$

With the same illumination intensity ($I_{con-ill} = I_{ls-ill}$) the ratio of the collected scattering power between two configurations is

$$\eta = \epsilon_{ls} / \epsilon_{con} = 0.61 \cdot \frac{NA_{con}^2}{NA_{ls-ill}}. \tag{6}$$

As mentioned, in our line-scanning setup, the illumination NA of our setup is $NA_{ls-ill} = 0.1$, while the detection NA is $NA_{ls-det} = 0.3$. Considering the confocal setup with same detection NA ($NA_{con} = 0.3$), the ratio of the collected scattering power between two configurations is $\eta = 0.549$ according to Equation (6). The dwell time is inversely proportional to the collected scattering signal $t_{ls}/t_{con} = 1/\eta \sim 1.8$.

Taken together, the expected theoretical value of $\alpha$ for our configuration is 16.39. This value is consistent with experimental values: Schlüßler et al.[3], who investigated zebrafish with a confocal configuration at our same wavelength, used laser power of 10 mW, i.e. equivalent laser intensity for NA=0.3 of $I_{con} = 5.06 \, mW/\mu m^2$ (note that in confocal configuration, the scattering efficiency is independent of NA) and the exposure time of $t_{con} = 200 \, ms$; this results into an increased efficiency (i.e. experimental value of $\alpha$) of 14.81 compared to our line-scanning geometry. It's important to notice, that the line-scanning configuration can be significantly improved in terms of geometrical efficiency by using focus extension technique; for example, by using the illumination NA of $NA_{ls-ill} = 0.6$ and a tunable lens to maintain the field-of-view, $\alpha$ is expected to be 2.7 (black line in **Figure 1f**), suggesting the light dose can be further reduced by about 5 times.

In summary, combining all contributions, the light dose delivered by our line-scanning configuration is 7.4%, i.e. more than one order of magnitude lower than an equivalent confocal configuration, while obtaining 10x speed advantage and 2x improved axial resolution.

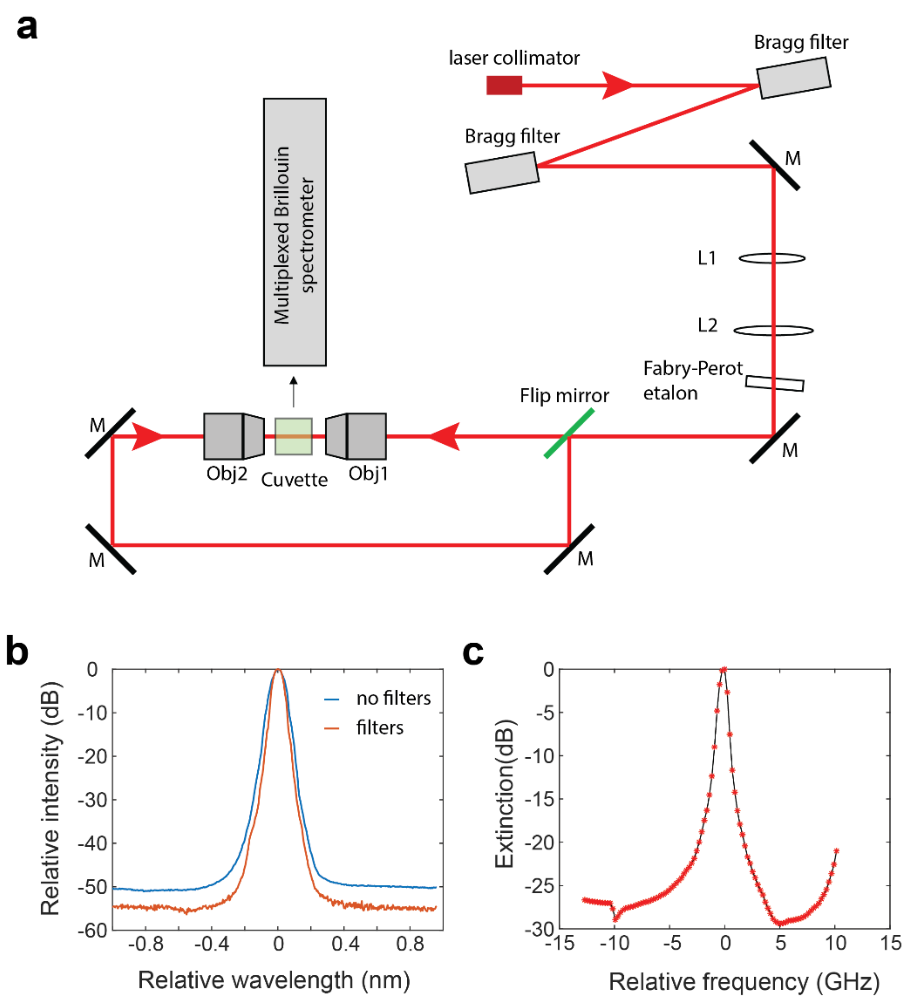


**Supplementary Fig.1**: Detailed schematic of the illumination beam. **a**, M, mirror; L1-L2, spherical lens with focal length of 20 mm and 80 mm, respectively; Obj1-Obj2, objective lenses (4x/0.1NA). **b**, The noise floor of the laser light is reduced by about 5 dB after passing through two Bragg filters. **c**, Extinction spectrum of the Fabry-Perot etalon used in the setup. For frequency component that is about 3.6 GHz (Brillouin shift of water) away from the laser frequency, the etalon can provide about 26 dB suppression. The value of zero in **b** and **c** represents the wavelength and frequency of the laser light, respectively.

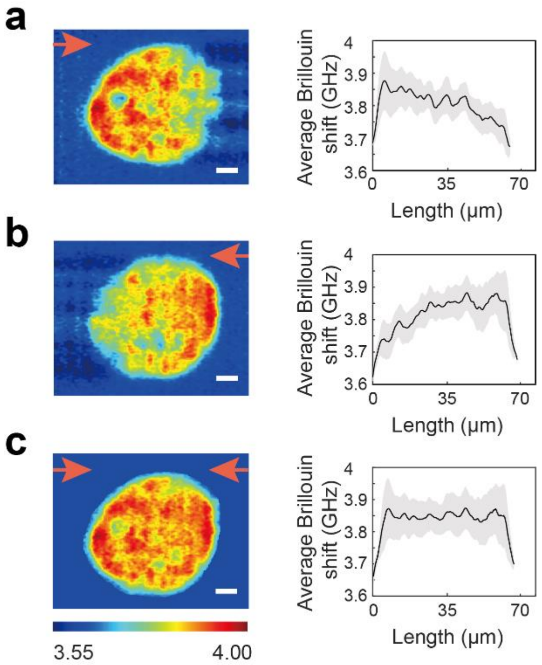


**Supplementary Fig.2**: Dual-line illumination. Brillouin image sections (left panels) and the averaged profile (right panels) along illumination direction of left-line illumination (**a**), right-line illumination (**b**), and combined dual-line illumination (**c**). The red arrow indicates the illumination direction of the beam line. Error bars represent the standard deviation. Scale bar, 10 μm. The color bar represents the Brillouin shift with the unit of GHz.

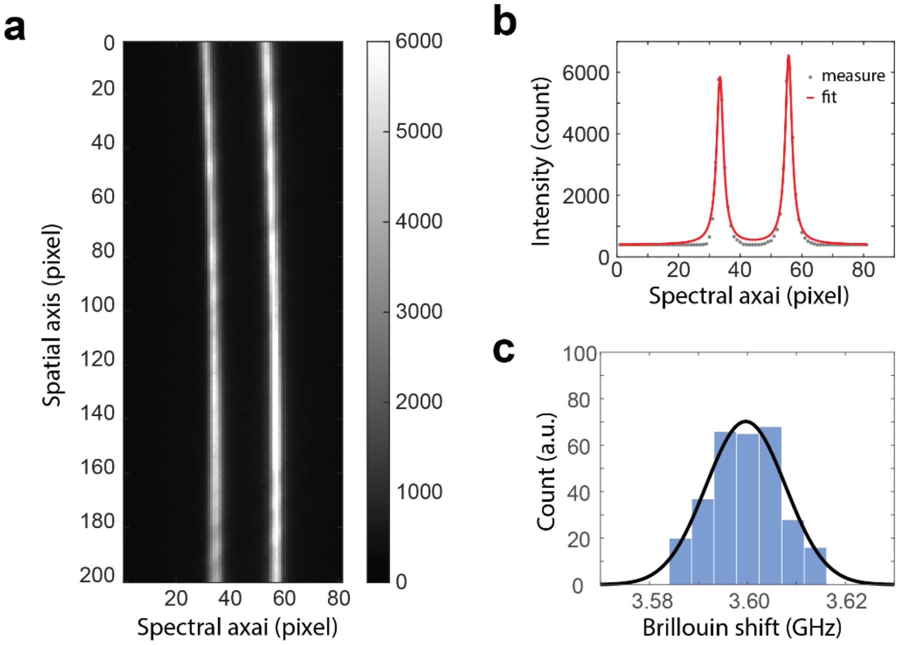


**Supplementary Fig.3**: Analysis of Brillouin spectrum. **a**, Examplary raw Brillouin spectrum of DI water acquired by the spectrometer of the LSBM setup with single-line illumination. **b**, Brillouin shift is extracted by fitting the spectrum of a single pixel with a Lorentzian function. Peaks represent the Stokes and anti-Stokes components of the Brillouin frequency. **c**, Histogram distribution of the estimated Brillouin shift of the same point after repeated acquisition (n=300). The standard deviation of the estimated Brillouin shift is 8 MHz.

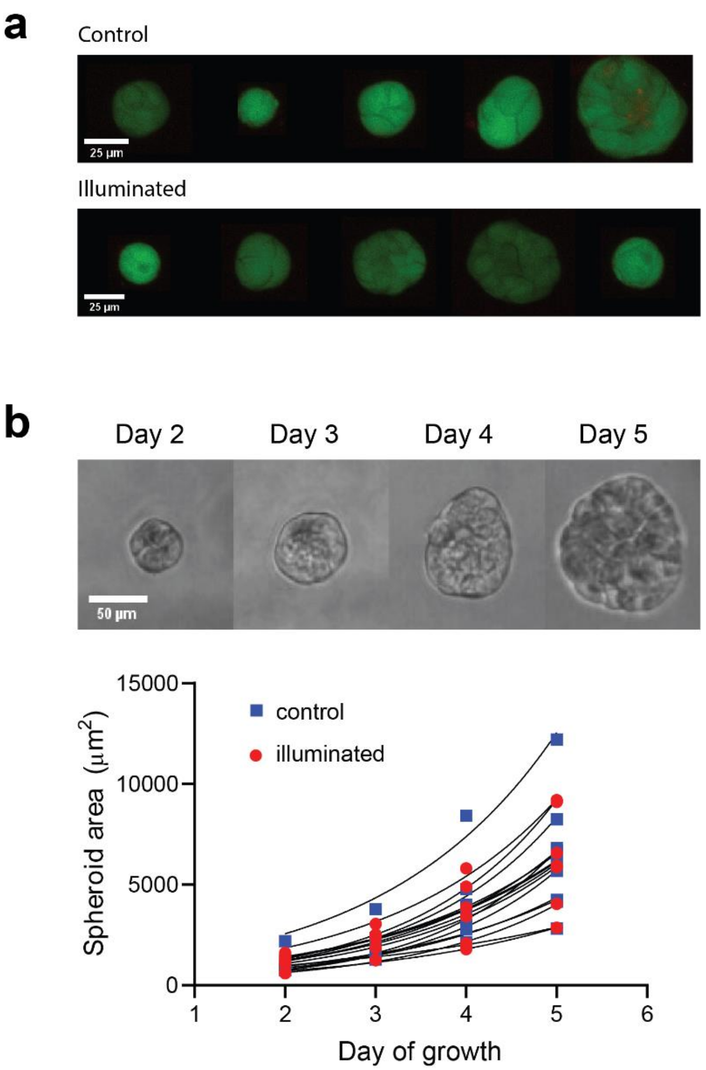


**Supplementary Fig.4**: Effect of light illumination on the viability and growth rate of the spheroids. **a**, Fluorescent images of the spheroids in control and illuminated groups. Live cells fluoresce in green (Calcein-AM) and dead cells fluoresce in red (EthD-1). **b**, Growth rate of the spheroids is not affected by laser illumination. The upper panel is the representative time-lapse image of a spheroid that is illuminated on Day 2. The points in the lower panel represent the area of spheroids over time. Black curves are best fit for exponential growth. Control group (n=10), Illuminated group (n=7).

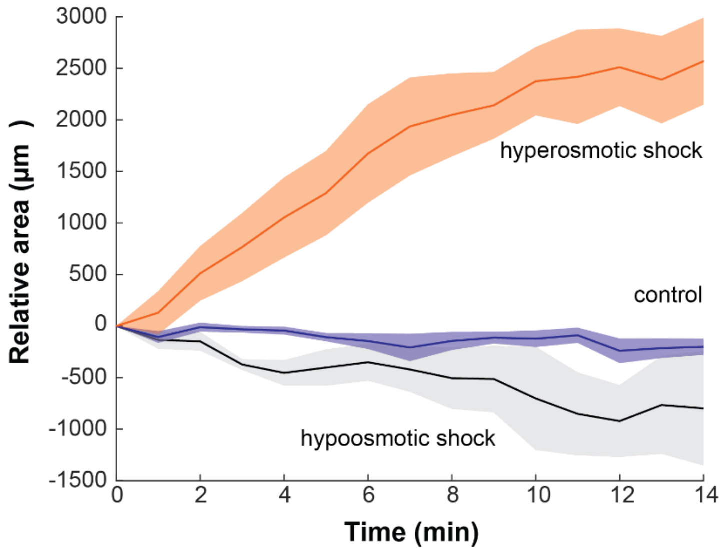


**Supplementary Fig.5**: Temporal change of the projection area of the spheroids under hyperosmotic shock (n=5), no shock (n=6), and hypoosmotic shock (n=5). Error bound indicates ± s.e.m..

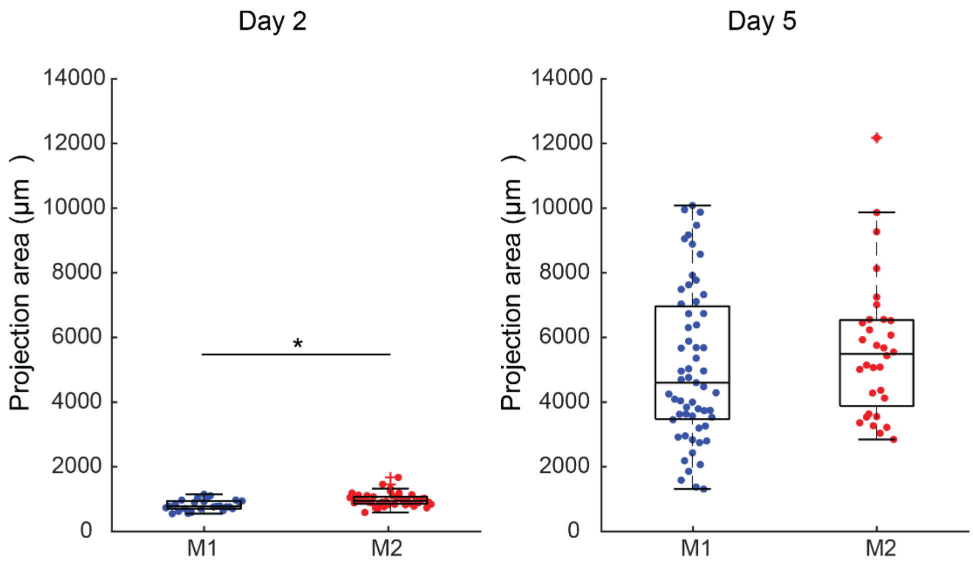


**Supplementary Fig.6**: Evolution of the spheroids' projection area. The averaged areas of M1 and M2 are 816 μm$^2$ and 978 μm$^2$ on Day 2 and increase to 5125 μm$^2$ and 5624 μm$^2$ on Day 5. **p*=0.007.

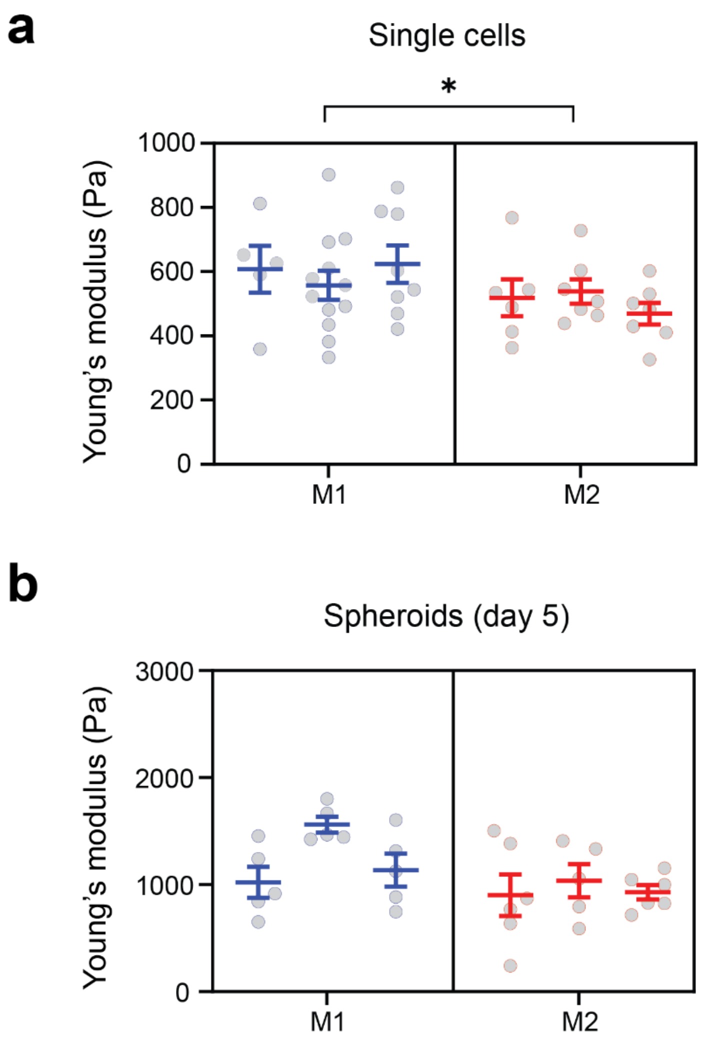


**Supplementary Fig.7**: All repeats of the Young's modulus measured by AFM. **a**, Indentation measurement on single cells. **b**, Indentation measurement on spheroids of Day 5. The mean ± s.e.m is used to represent each repeat. Statistical significance is determined by performing unpaired t-test with all repeats. * $p$=0.038.
